# Laser-driven Wireless Deep Brain Stimulation using Temporal Interference and Organic Electrolytic Photocapacitors

**DOI:** 10.1101/2021.11.02.466993

**Authors:** Florian Missey, Mary J. Donahue, Pascal Weber, Ibrahima Ngom, Emma Acerbo, Boris Botzanowski, Ludovico Migliaccio, Viktor Jirsa, Eric Daniel Głowacki, Adam Williamson

## Abstract

Deep brain stimulation (DBS) is a technique commonly used both in clinical and fundamental neurosciences. Classically, brain stimulation requires an implanted and wired electrode system to deliver stimulation directly to the target area. Although techniques such as temporal interference (TI) can provide stimulation at depth without involving any implanted electrodes, these methods still rely on a wired apparatus which limits free movement. Herein we report organic photocapacitors as untethered light-driven electrodes which convert deep-red light into electric current. Pairs of these ultrathin devices can be driven using lasers at two different frequencies to deliver stimulation at depth via temporally interfering fields. We validate this concept of laser TI stimulation using numerical modeling, *ex vivo* tests with phantom samples, and finally *in vivo* tests. Wireless organic photocapacitors are placed on the cortex and elicit stimulation in the hippocampus, while not delivering off-target stimulation in the cortex. This laser-driven wireless TI evoked a neuronal response at depth that is comparable to control experiments induced with deep brain stimulation protocols using implanted electrodes. Our work shows that a combination of these two techniques – temporal interference and organic electrolytic photocapacitors – provides a reliable way to target brain structures requiring neither deeply implanted electrodes nor tethered stimulator devices. The laser TI protocol demonstrated here address two of the most important drawbacks in the field of deep brain stimulation and thus holds potential to solve many issues in freely-moving animal experiments or for clinical chronic therapy application.

## INTRODUCTION

Targeted electrical stimulation in neuroscientific research is an invaluable tool, as well as the methodology of choice in an increasing range of clinical applications. Indeed, experimental research uses electrical brain stimulation in numerous ways, for example mapping the sensory motor corticies^1,2,3^ or delineating epileptogenic zones^4,5^ in patients. However, a main drawback is that in the majority of applications the electrical stimulation is most commonly delivered via invasive wired electrodes. In contrast, non-invasive neuromodulation without an implanted device in the brain can be achieved via transcranial direct current stimulation (tDCS)^6^, transcranial focused ultrasound stimulation^7^, or transcranial magnetic stimulation (TMS)^8^. Although these non-invasive procedures show great promise stimulating cortical structures, focally reaching deep brain structures is still a significant challenge. Furthermore, the cited non-invasive stimulation methods require complex and cumbersome equipment making adoption for freely-moving animal studies complicated. Recently, a non-invasive but wired deep brain stimulation was introduced, in mice, by Grossman et al.^9^: temporal interference (TI) stimulation. This stimulation method relies on frequencies higher than 1 kHz. Such frequencies do not elicit a response from neurons and propagate through tissue with relatively low attenuation. When two carrier frequencies are used with a frequency offset, interference patterns at the offset frequency can be focally observed. This lower frequency envelope has been shown experimentally to achieve focal stimulation where the two carrier frequencies maximally constructively interfere. The exact mechanisms of stimulation using TI envelopes at the cellular level are a current topic in contemporary research^10^.

We recently demonstrated successful non-invasive TI stimulation, specifically orientation-tunable TI to create focal hippocampal stimulation in mice.^11^ However, this type of TI stimulation still requires a not negligeable number of wires and complex equipment and thus is limited in its range of applications. In the present work we show that our previously demonstrated TI protocol^11^ can be applied wirelessly by using ultrathin, laser-driven photocapacitors. First reported in a series of papers^12,13^ in 2018, organic electrolytic photocapacitors (OEPCs) can convert deep-red light illumination into electrical impulses at biological interfaces (OEPC function detailed in supplementary figure 3)^14^. OEPCs are fully wireless and are essentially floating stimulators, making them potentially ideal for TI. Herein, we utilize the OEPC, with an additional conductive polymer coating, Using PEDOT:PSS, which has been demonstrated to improve photostimulation efficiency^15^, for a minimally invasive wireless deep brain stimulation protocol. Using PEDOT:PSS on the OEPC greatly increases the charge storage capacity^16^ and thus enhances the stimulation for the same illumination.

To recreate our TI stimulation protocol for targeting the hippocampus, we placed two OEPCs on top of the *dura* matter of mice cortex to act as stimulation electrodes (Fig. 1). As explained in Figure 1, each OEPC is driven at either 1250Hz or 1300Hz using deep-red (638nm) light diodes, where kHz light modulation is translated into kHz electrical brain stimulation. This approach leads to an envelope at the offset of the carrier frequencies (50 Hz), analogous to standard TI and creating a stimulation hotspot at depth in the tissue. We demonstrate evoked epileptiform activity, more precisely Inter-Ictal-Like Event (IILEs), directly following laser TI stimulation in the CA3 to CA1 region of the mice hippocampus (Schaffer collaterals). Standard TI protocols can stimulate deep brain regions without any implantable device directly in the brain, thus avoiding physical damage of neural tissue. However, all the regular protocols require a wired set up to carry out the stimulation. Our Laser TI protocol has the advantage of being wireless and portable. This optical, wireless method could provide an interesting advantage for certain *in vivo* animal experimental setups, as well as great potential for clinical investigation/therapy.

**Figure 1.**
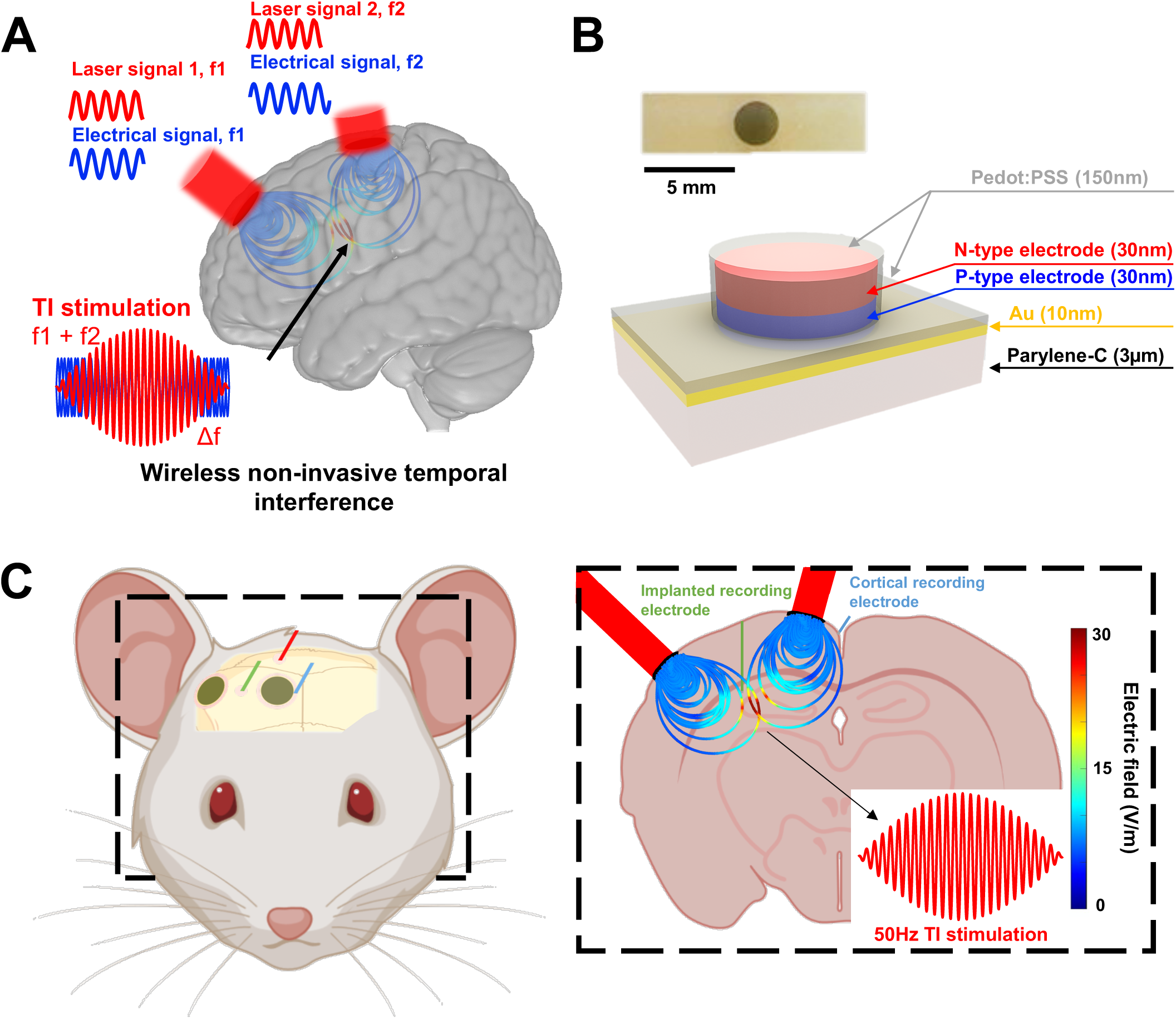
Laser-driven Temporally Interfering fields (Laser TI). **A**, Temporal interfering fields are usually realized using a wired apparatus^6^. The combination of OEPC^12,13,14,15,17^ technology with TI protocols provides a minimally invasive and wireless deep brain stimulation possibility. Analogous to classical TI protocols, field lines are generated using deep red light illumination (638nm) and interfere at the targeted deep brain region, the CA3 to CA1 junction in the hippocampus. B, The OEPC devices (both cross-sectional schematic and top-view photograph shown) are made on ultrathin plastic substrates of parylene-C, modified with a conducting gold layer. Circular PN pixels constitute the primary stimulation photoelectrodes. Both the front and back electrodes are modified with conducting polymer PEDOT:PSS in order to lower impedance.

## RESULTS

### Characterization of optimal OEPC placement

We developed a custom implantation protocol to target the CA3 to CA1 junction of the hippocampus (Schaffer collaterals). To find the optimal location for OEPC placement, a finite element model was created with a 3D mouse brain and two OEPCs to scale (Fig. 2A). Simulations were run corresponding to the *in vivo* experiment, namely 1250Hz and 1300Hz as carrier frequencies and 10 s of stimulation. Using a FEM model, optimal placement of the OEPCs could be found to induce a maximal Laser TI stimulation at the hotspot and minimal stimulation in the cortex. Electric field distribution and associated recordings with envelope calculation in the hippocampus or in the cortex are shown in Figure 2.

**Figure 2.**
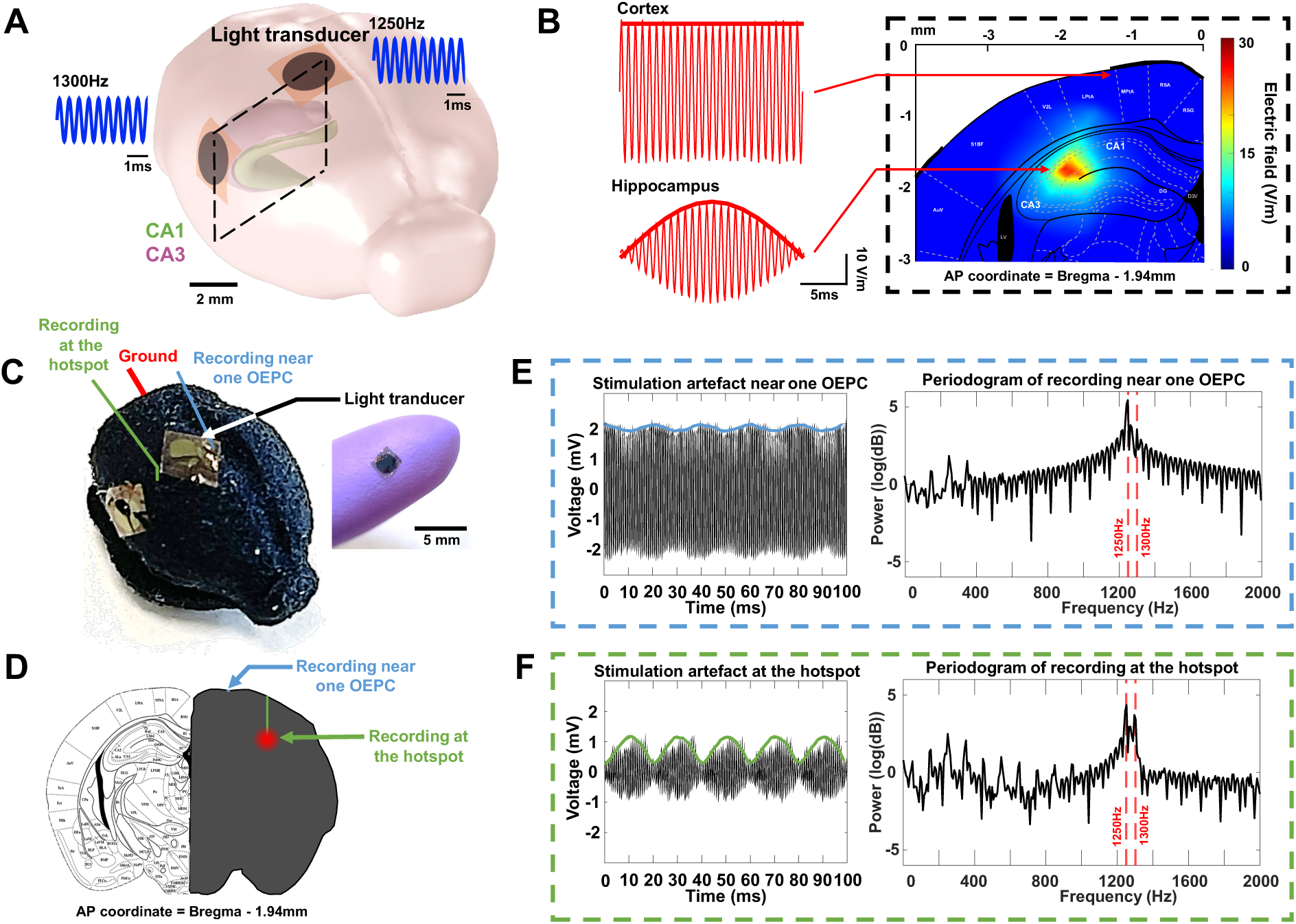
*Ex vivo* validation of Laser TI. Temporal interference protocols are highly dependent on an optimal positioning of the stimulating electrodes. (**A** and **B**) Using a Finite Element Method (FEM) model and implementing both OEPC geometries and electrophysical properties of the brain and OEPCs (Fig. S1), we could recreate an artificial Laser TI system and optimize OEPC placement (**B**). In addition to the placement optimization, the FEM provided *in silico* evidence for TI using these devices. (**C**) Laser TI was also tested on a mouse phantom gel with equivalent proportions and the OEPCs were placed based on results from the FEM model. An OEPC device is shown laminated onto a gloved fingertip for scale and (D) the utilized coordinates are displayed according to Paxinos mouse brain atlas (AP = Bregma - 1.94mm). Local field potentials were recorded near one OEPC (**E**) and at the hotspot (**F**, hippocampus location) to illustrate the temporal interference stimulation.

### Laser TI artefact recording in the hippocampus and in the cortex

To ensure that the *in silico* and *ex vivo* phantom results could be recreated *in vivo*, we recorded local field potentials (LFPs) from both recording electrodes (hippocampus and cortex) during the light stimulation (Fig. 3). As expected, the stimulation artefact recorded in the hippocampus is well defined with a clear amplitude-modulated 50 Hz envelope, while the signal in the cortex is less pronounced with amplitude modulation that is more difficult to distinguish (Fig. 3). When analyzing the frequency spectra of these artefacts via periodograms, the LFPs in the hippocampus show two peaks at 1250Hz and 1300Hz with the same amplitude; thus demonstrating the optimal frequency combination in the hippocampus. In the periodogram for the cortical location, we show that the 1250Hz peak is weaker than the 1300Hz, meaning that the combination of the frequencies was not as strong, as desired. This difference in the frequency combination, essentially the amount of amplitude modulation in the sine wave signal, at various locations is expected based on FEM modeling and explains the differences in the artefact amplitudes.

**Figure 3.**
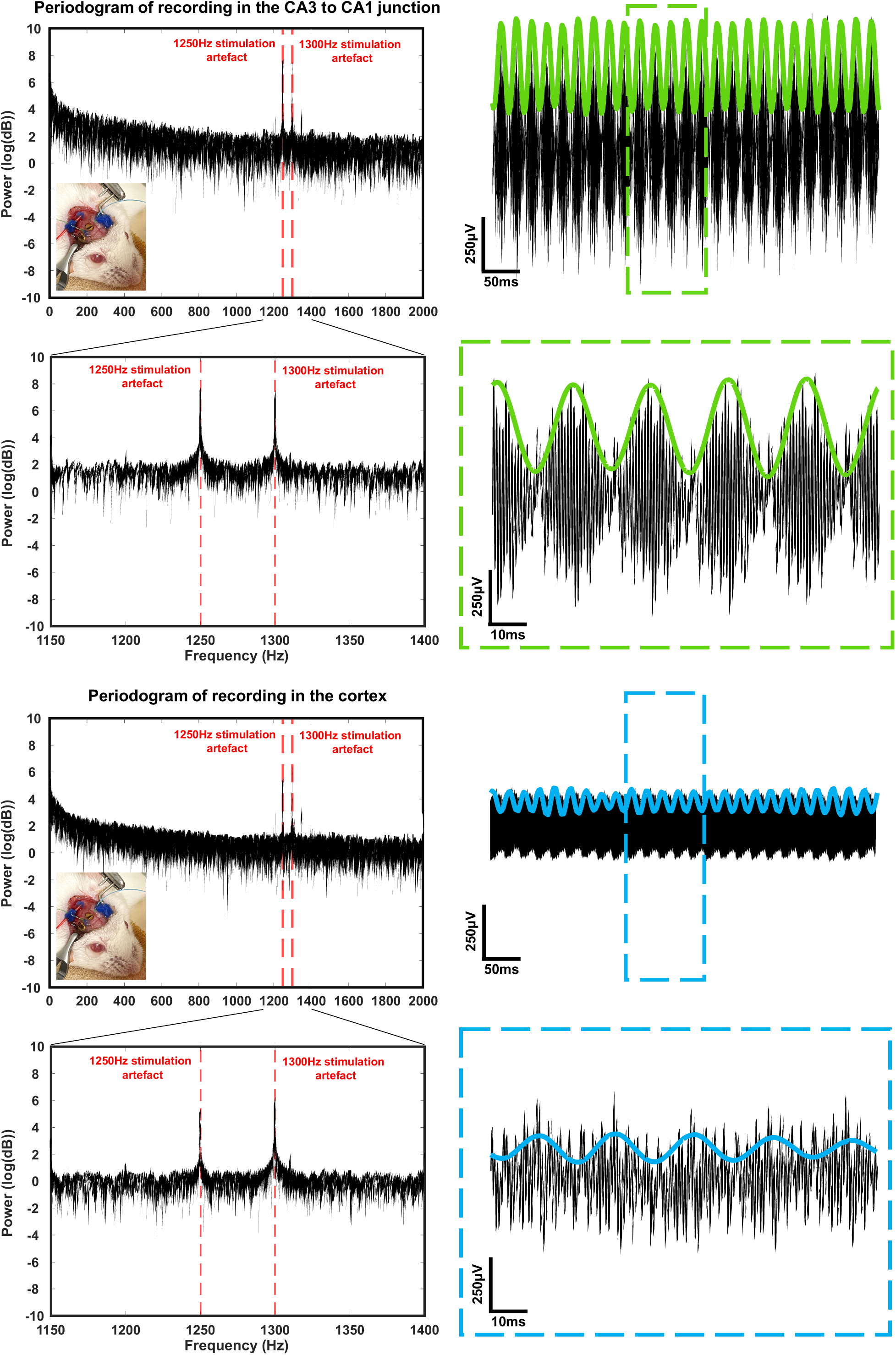
Temporal interference artefacts during Laser TI stimulation. Recordings during stimulation show clear Laser TI in the hippocampus. By selecting 1300 Hz and 1250 Hz as our stimulation frequencies, we could elicit a 50 Hz stimulation envelope at depth in the mouse hippocampus. Both raw recordings and frequency component analysis show that the frequency mixing is ideal in the hippocampus and less pronounced in the cortex, near one stimulation OEPC.

### Evoked IILE using Laser TI stimulation

In order to characterize the IILEs evoked with the Laser TI stimulation and to compare them with the those elicited using a standard stimulation protocol, we analyzed the frequency components, the duration, and the amplitude of all events. IILEs evoked in both protocols are similar when observing raw LFP or spectrograms (Fig. 4A and 4B). For a more detailed analysis, a mean periodogram was realized by gathering all IILEs evoked in each group. As expected, evoked epileptiform activity has an increase of the frequency component in the beta/gamma band which is characteristic of epileptic tissue (Fig. 4C). When looking at the durations and amplitudes of each IILE, we could not find any differences in between the two groups.

**Figure 4.**
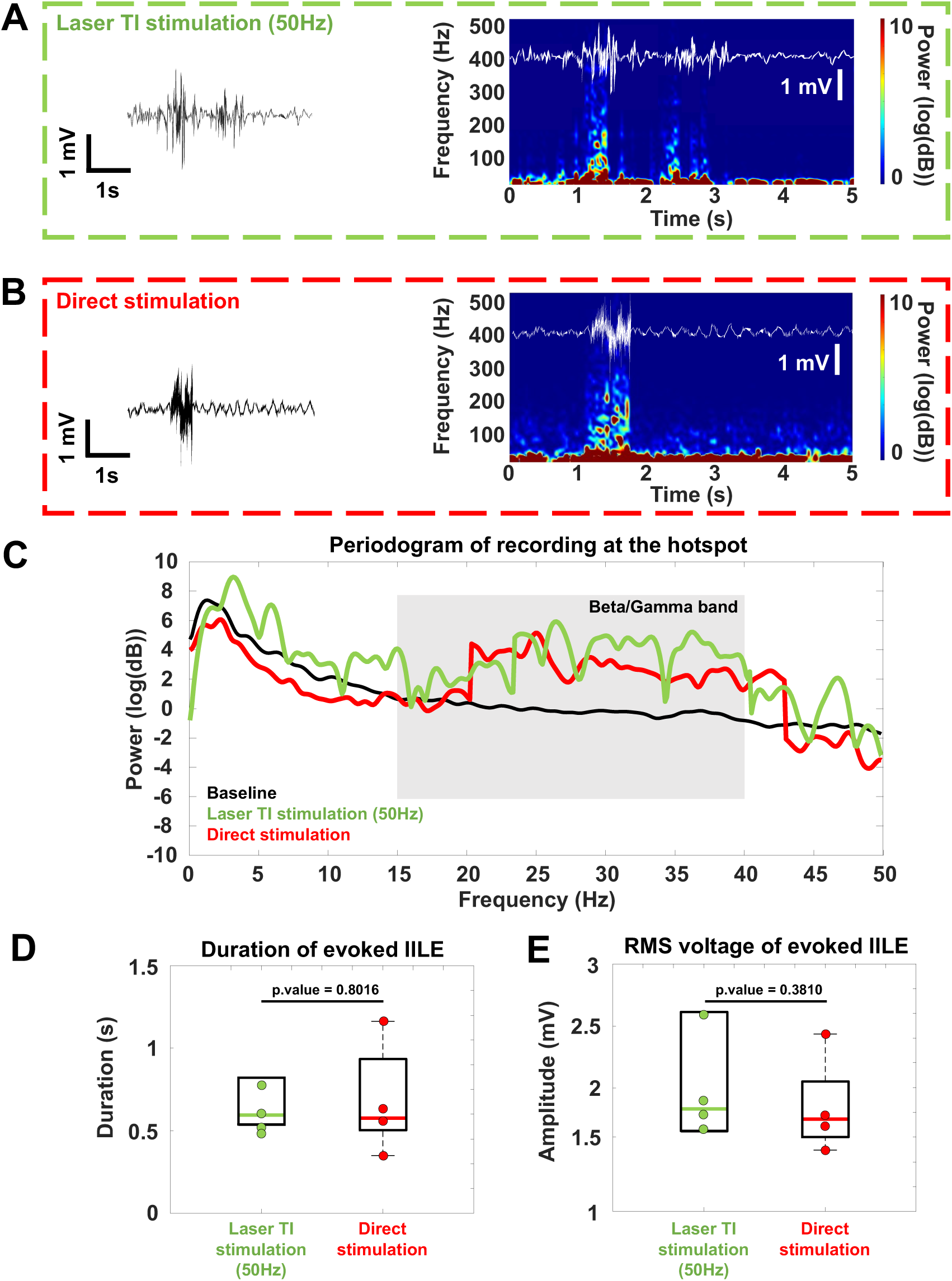
Evoked epileptiform activities after Laser TI stimulation. Electrophysiological epileptiform activities were evoked after all Laser TI stimulations. An example of the evoked IILE is depicted in **A** and **B**. IILE were analyzed in the time (**A**,**B**)and frequency domains (**C**) using temporal plot or spectrograms to illustrate both the epileptiform shape of the activity and the increase in beta/gamma range. IILEs were gathered and an average periodogram was plotted. Post Laser TI stimulation (n=4) IILEs were compared to regular IILE evoked with direct electric stimulation (n=4) of the hippocampus, no differences were showed (**D, E**).

## DISCUSSION

Central nervous system stimulation typically relies on implanted electrodes that can directly provide electrical stimuli to brain tissue. However, researchers and clinicians continually face the dilemma between using an invasive technique with high spatial focality^18,19,20^ but damaging the tissue, or a non-invasive method with a shallower and wider stimulation^21,22,23,24^ which avoids risk for the brain. Temporal Interference has been demonstrated as a new stimulation technique can overcome these issues by allowing a deep and focal stimulation using only minimally invasive brain implants^9,11^. Exploiting the intrinsic property of neurons, that they are not stimulated by frequencies above 1 kHz^25^, temporal interferences stimulation uses two distinct frequencies above this 1 kHz threshold that will meet at a target depth and create a stimulation envelope with a desired frequency^11^. Although this technique is very promising in term of minimally invasive deep brain stimulation, it requires a great deal of wiring to the appropriate tools for signal delivery. Engineering achievements in micro-optoelectronic design present an alternative as light can be converted into useful electric signals^12,13,14,15^. The aim of OEPCs, which fall into this category, is to produce an electrical stimulus using deep red light, thus enabling wireless electrical stimulation. The combinations of minimally invasive OEPCs with temporal interference stimulation protocols allows for wireless deep brain stimulation using only two LED sources. In this work we suggest that the use of laser-driven TI can be an alternative to other stimulation techniques for the central nervous system. Indeed, when triggering two different OEPCs with two different current sources, each of them will transduce their own light signal into electric fields that will interfere, creating a hotspot at the desired target area. This way laser-driven TI can focally activate deep brain structures with a high resolution and improve regular TI stimulation by making it totally wireless. The laser-driven TI can generate comparable electrophysiological patterns as those demonstrated in previous direct electrical stimulation experiments^26,27^.

In summary, laser TI stimulation can be used as an alternative to electrical TI stimulation or direct electrical stimulation. The ability of the OEPCs to be optimally activated by deep red-light LED make laser TI an advantageous portable option for central nervous system stimulation. Deep red light is known to penetrate through biological tissues with minimal attenuation^28^. This makes the use of OEPCs a great opportunity for the introduction of laser TI in chronic deep brain stimulation protocols. The electrical signal generated by the OEPCs is directly proportional to the light intensity sent by the laser, enabling a fine control of the electrical signal transmitted to the tissue. Finally, as a therapeutic tool in the clinic, laser TI can be used to stimulate typical brain regions as well as regions that are not able to be stimulated with standard implantation protocols. Thus, Laser TI can provide a wireless, deep, and focal brain stimulation with a large panel of possible targets that can be integrated in all deep brain stimulation protocols.

## MATERIALS AND METHODS

### Animals

Animal experiments were performed under the agreement of European Council Directive EU2010/63 and French Ethics approval (comité d’éthique en experimentation animale n°70 - Williamson, n. APAFIS 20359 - 2019041816357133). 8 (2 groups of 4 mice) OF1 (Oncins France 1, Charles Rivers Laboratories, France) aged 8 to 10 weeks old underwent the surgery and stimulation protocol described below. Mice were housed in cage of 4 and under a normal 12/12h day-night cycle at room temperature, water and food were dispensed *ad libitum*.

### Surgical procedure

Mice were anesthetized using a xylazine (20mg/kg) and ketamine (50mg/kg) mix via an intraperitoneal injection (2.5µL/g). During the surgeries, the mice temperature was monitored, and their eyes were covered with vitamin A to avoid any damages. A wide incision was made in the skin and a modified metal clip was cemented on the skull (cement primer Ultradent® and photopolymerized resin Lc Block-Out GACD®) to be used as a stabilizer for the mouse head. For the Laser TI group (n=4), two craniotomies of about 3mm diameter were realized with the following coordinates (AP = Bregma −1.94 / ML_center_ = Bregma + 0.5 and Bregma + 4.5), taking care to not damage the *Dura mater*. OEPCs (3mm diameter) were gently placed on the craniotomy windows. Two tungsten electrodes (70µm diameter) were inserted, one in the Cornu Ammonis 3 (CA3) to Cornu Ammonis 1 (CA1) junction in the hippocampus (AP = Bregma −1.94 / ML = Bregma + 2.25 / DV = −1.27) and the other one near one OEPC. For the control group (n=4) that will receive direct electric hippocampal stimulation, a twisted pair deep brain stimulation electrode (2×125µm, platinum) was inserted at the following coordinates (AP = Bregma −1.94 / ML = Bregma + 2.8 / DV = −1.57 / Θ= 20°). Finally, and for both groups, a mini pin was implanted in the cerebellum to serve as ground and reference.

### Light stimulation and recording

OEPCs modified with PEDOT:PSS were fabricated according to previously reported procedures^8,19^. First a 2.2 µm thick parylene-C substrate was deposited on a glass carrier wafer^8,19^. The parylene-C surface was then activated with O_2_ plasma (Diener electronic GmbH, 50 W), and vapor-phase deposition of 3-(mercaptopropyl)trimethoxysilane was carried out to enhance gold adhesion. A 10 nm thick, semi-transparent Au layer was then deposited by thermal evaporation onto the treated parylene-C substrate. This layer acts as the return electrode of the final device. Subsequently, the organic PN photoelectrode was deposited via thermal evaporation through a shadow mask, defining pixels of 3 mm in diameter. The PN junction consisted of H2Pc, (Phthalocyanine, 30 nm, Alfa Aesar) and PTCDI (N,N’-dimethyl-19 3,4,9,10-perylenetetracarboxylic diimide, 30 nm, BASF), which prior to deposition were purified by threefold temperature-gradient sublimation. Upon illumination, photogenerated electrons accumulate at the PTCDI / electrolyte interface while holes are driven to the Au return electrode / electrolyte interface. Finally, a PEDOT:PSS (CLEVIOS PH 1000) formulation including 2% w/w (3-glycidyloxypropyl)trimethoxysilane (GOPS) was spin-coated at 1500 rpm using a 1000 rpm.s^-1^ acceleration for 60 s to obtain a well-adhered PEDOT:PSS coating (150 nm ± 5 nm, measured using scanning stylus profilometry). Following fabrication, devices were cut to size using a scalpel and then delaminated from the underlying glass carrier wafer. Once delaminated, the OEPC devices were manipulated using tweezers and could be adhered to the desired region of the exposed dura matter. For all light stimulation sessions, the mice eyes were covered with aluminum foil. We used a Four-Channel LED driver 10A coupled with a Keysight® EDU33212A in order to generate light pulses with custom intensity and frequency. Two 1.2W 638nm laser diodes were triggered respectively with 1250 Hz and 1300 Hz. Before the light stimulation protocol, the laser intensity was set to 5% to visually target the OEPCs location safely. A baseline neural recording using the implanted tungsten electrodes and an Intan 128ch Stimulation/Recording Controller (IntanTech®) was also conducted before any light stimulation. The light stimulation protocol consists of a 10-second stimulation train with both diodes set to 50% (1200W, max current 1000mA). One OEPC is triggered with a 1250Hz stimulus and the other with a 1300Hz stimulus. It should be noted that sine waves for the laser TI protocol have a DC offset to ensure the LED is only driven with positive values. Simultaneously, Local Field Potentials (LFPs) were recorded with the two tungsten electrodes placed in the hippocampus and near one OEPC.

### Direct electric stimulation

A standard hippocampus stimulation was realized using the Intan 128ch Stimulation/Recording Controller. The protocol consists in a 10 second bipolar stimulation with biphasic pulses of 1ms (50% duty cycle) sent at 50Hz. Current amplitudes of stimulation were increased until finding IILE in the target area.

### Numerical simulation

A 3D mouse brain model designed in Blender® was loaded in COMSOL Multiphysics® software, version 5.5 (www.comsol.com) to perform finite-element simulations. OEPC representations were placed on top of the 3D mouse brain and physical properties such as permittivity and conductivity were allocated to both device and biological models. Electric current was assigned to each OEPC representation with the respective frequencies used in the *in vivo* protocol (1250 and 1300Hz). Resulting voltages and field lines were then investigated through 3D distribution plots directly in COMSOL. For a better comparison with the *in vivo* results, the simulated voltage recorded at the hotspot was extracted and placed in a MATLAB (MathWorks) matrix.

### Statistical analysis

*In silico* simulations or *in vivo* LFP recordings were plotted and analyzed using MATLAB (MathWorks). All resulting envelope functions were also calculated with MATLAB (MathWorks) using a Hilbert function. Inter-Ictal-Like Events (IILEs) were defined as activity with an amplitude of 2 × baseline. The duration and amplitudes (RMS of the voltage over the IILE duration window) were gathered and statistics were performed using R software. Normality Shapiro tests were realized on both groups (Laser TI and Direct stimulation) and non-parametric tests (Wilcoxon-Mann-Whitney and Friedman, power = 0.8) were performed to bring out any differences between these two groups.

## ACKNOWLEDGMENTS

A.W. and E.D.G. acknowledge funding from the European Research Council (ERC) under the European Union’s Horizon 2020 research and innovation programme (A.W. grant agreement No 716867; E.D.G. grant agreement No. 949191). E.D.G. gratefully acknowledges financial support from the Knut and Alice Wallenberg Foundation within the framework of the Wallenberg Centre for Molecular Medicine at Linköping University and the Swedish Research Council (Vetenskapsrådet, 2018-04505).

## AUTHOR CONTRIBUTIONS

A.W. conceived the project. F.M. and P.W. performed experiments. F.M. ran finite-element simulations and analyzed neural data. E.D.G., M.J.D., L.M. and A.W. designed organic photocapacitors. F.M., A.W., and E.D.G. wrote the paper with input from the other authors including M.J.D., I.N., E.A., B.B. and V.J.

## SUPPLEMENTARY FIGURES

**Figure S1.**
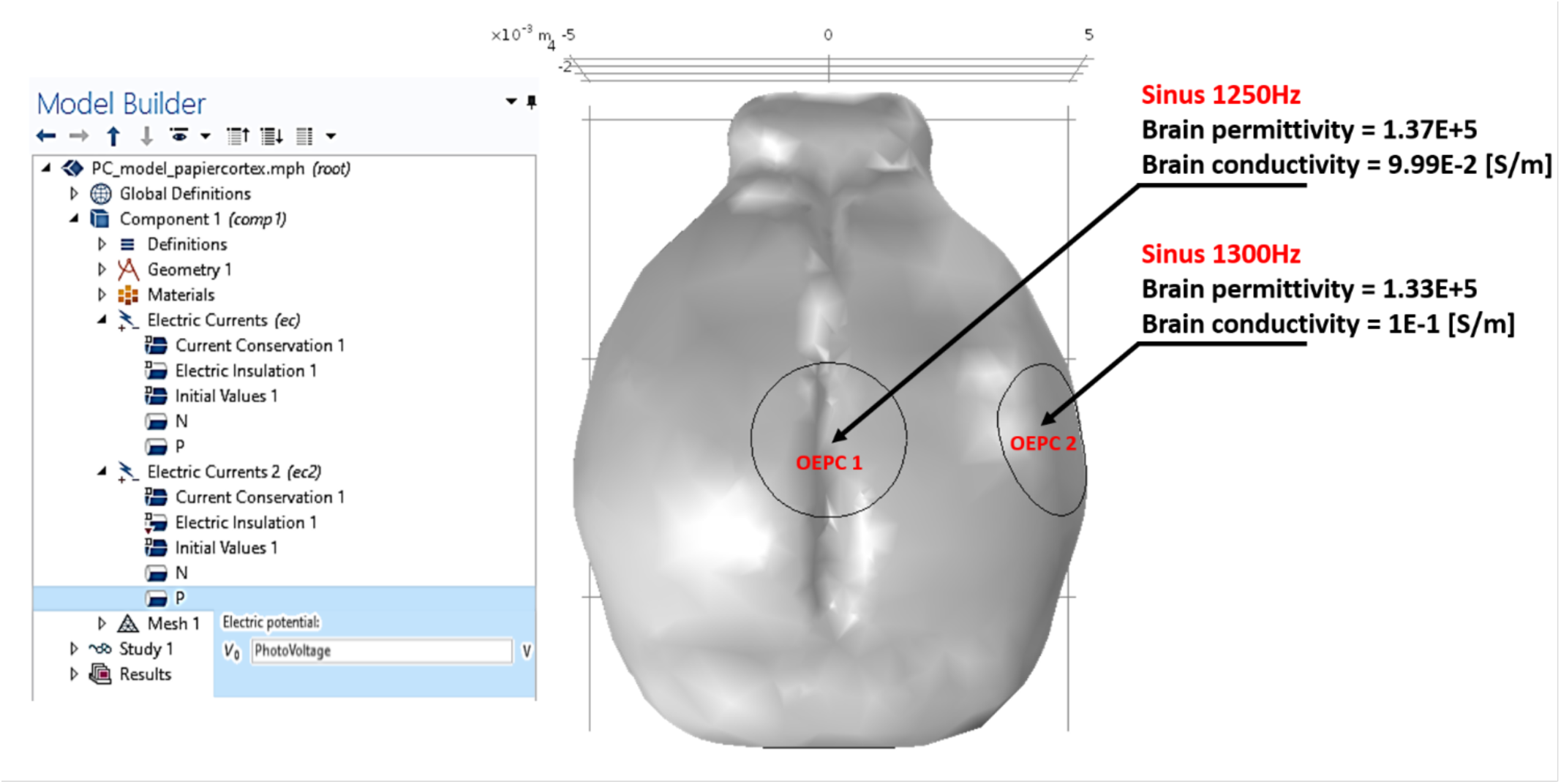
Finite Element Model and custom parameters for Laser-driven wireless temporal interference.

**Figure S2.**
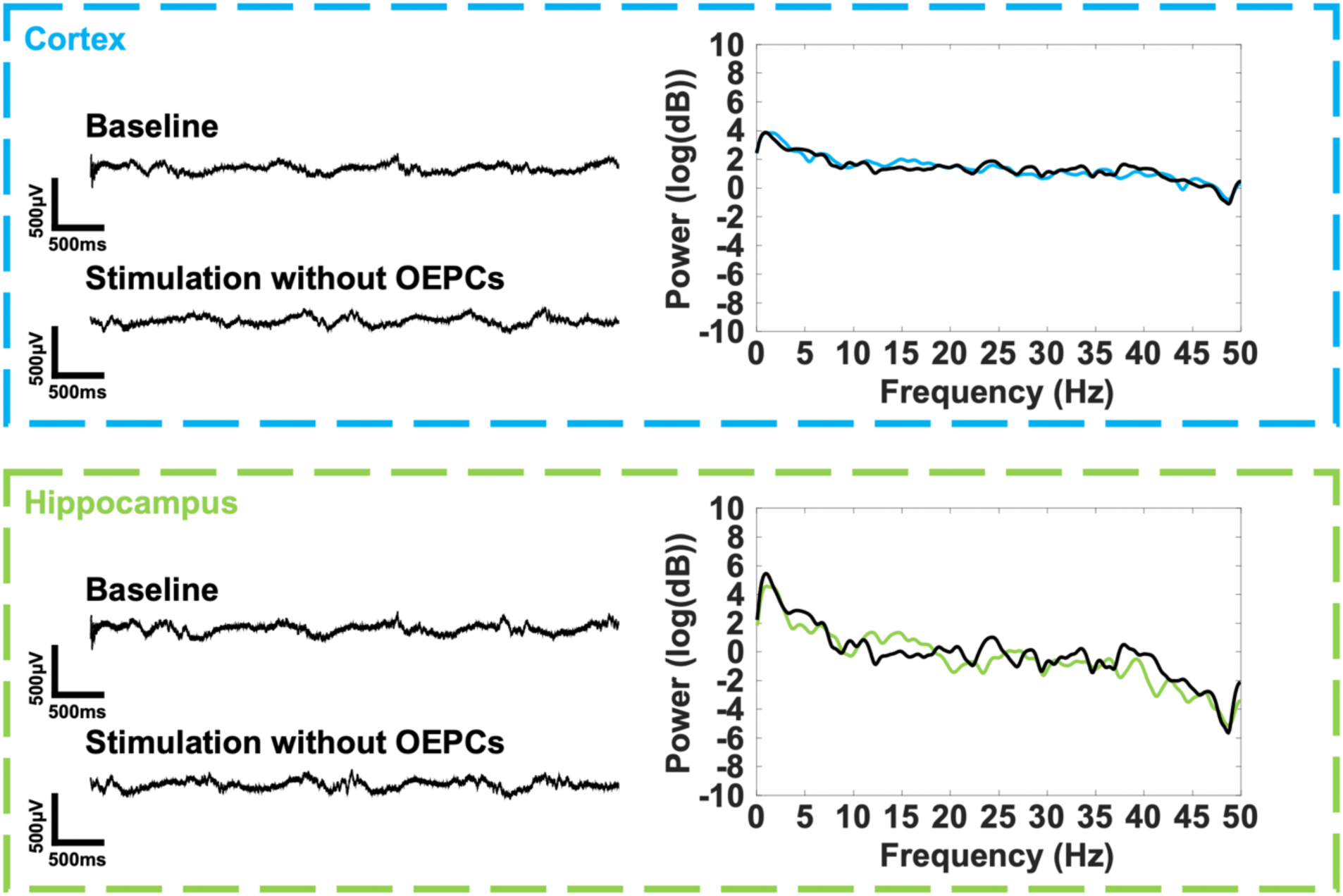
Non-evoked activity following light stimulation with no OEPCs.

**Figure S3.**
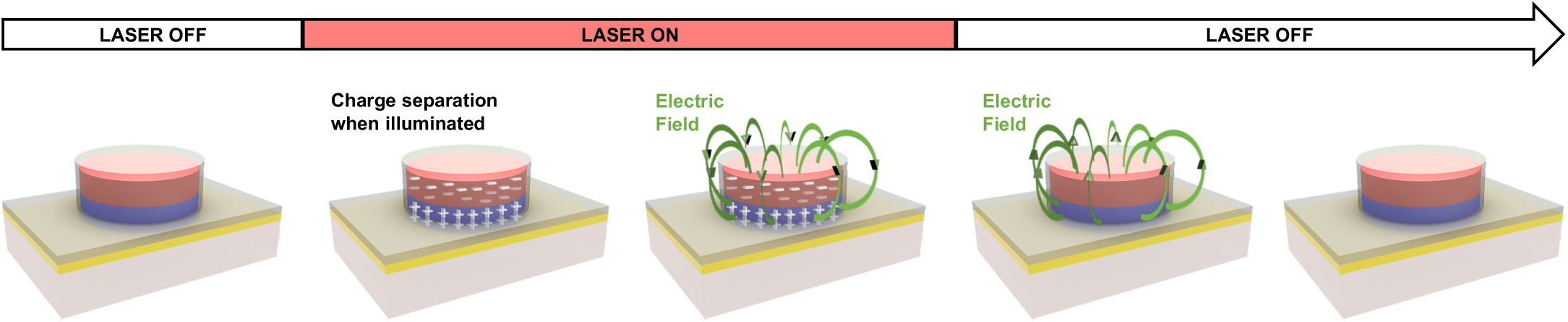
Light conversion into electric field using OEPCs.

## REFERENCES

1. Penfield, W. & Boldrey, E. SOMATIC MOTOR AND SENSORY REPRESENTATION IN THE CEREBRAL CORTEX OF MAN AS STUDIED BY ELECTRICAL STIMULATION. Brain 60, 389–443 (1937).

2. Borchers, S., Himmelbach, M., Logothetis, N. & Karnath, H.-O. Direct electrical stimulation of human cortex — the gold standard for mapping brain functions? Nat Rev Neurosci 13, 63–70 (2012).

3. Ritaccio, A. L., Brunner, P. & Schalk, G. Electrical Stimulation Mapping of the Brain: Basic Principles and Emerging Alternatives. Journal of Clinical Neurophysiology 35, 86–97 (2018).

4. George, D. D., Ojemann, S. G., Drees, C. & Thompson, J. A. Stimulation Mapping Using Stereoelectroencephalography: Current and Future Directions. Front. Neurol. 11, 320 (2020).

5. Jayakar, P. et al. Diagnostic utility of invasive EEG for epilepsy surgery: Indications, modalities, and techniques. Epilepsia 57, 1735–1747 (2016).

6. Das, S., Holland, P., Frens, M. A. & Donchin, O. Impact of Transcranial Direct Current Stimulation (tDCS) on Neuronal Functions. Front. Neurosci. 10, (2016).

7. Lee, W. et al. Transcranial focused ultrasound stimulation of human primary visual cortex. Sci Rep 6, 34026 (2016).

8. Hallett, M. Transcranial magnetic stimulation and the human brain. Nature 406, 147–150 (2000).

9. Grossman, N. et al. Noninvasive Deep Brain Stimulation via Temporally Interfering Electric Fields. Cell 169, 1029-1041.e16 (2017).

10. Mirzakhalili et al., Biophysics of Temporal Interference Stimulation. Cell Systems 11, 557–572 (2020).

11. Missey, F. et al. Orientation of Temporal Interference for Non-invasive Deep Brain Stimulation in Epilepsy. Front. Neurosci. 15, 633988 (2021).

12. Rand, D. et al. Direct Electrical Neurostimulation with Organic Pigment Photocapacitors. Adv. Mater. 30, 1707292 (2018).

13. Jakešová, M. et al. Optoelectronic control of single cells using organic photocapacitors. Sci. Adv. 5, eaav5265 (2019).

14. Missey, F. et al. Organic electrolytic photocapacitors for stimulation of the mouse somatosensory cortex. http://biorxiv.org/lookup/doi/10.1101/2021.10.20.465090 (2021) xdoi:10.1101/2021.10.20.465090.

15. Silverå Ejneby, M. et al. Extracellular Photovoltage Clamp Using Conducting Polymer-Modified Organic Photocapacitors. Adv. Mater. Technol. 5, 1900860 (2020).

16. Donahue, M. J. et al. Tailoring PEDOT properties for applications in bioelectronics. Materials Science and Engineering: R: Reports 140, 100546 (2020).

17. Silverå-Ejneby, M. et al. A chronic photocapacitor implant for noninvasive neurostimulation with deep red light. http://biorxiv.org/lookup/doi/10.1101/2020.07.01.182113 (2020) doi:10.1101/2020.07.01.182113.

18. Jeffrey, M. et al. A reliable method for intracranial electrode implantation and chronic electrical stimulation in the mouse brain. BMC Neurosci 14, 82 (2013).

19. Mullin, J. P. et al. Is SEEG safe? A systematic review and meta-analysis of stereo-electroencephalography-related complications. Epilepsia 57, 386–401 (2016).

20. Rossini, P. M. et al. Non-invasive electrical and magnetic stimulation of the brain, spinal cord, roots and peripheral nerves: Basic principles and procedures for routine clinical and research application. An updated report from an I.F.C.N. Committee. Clinical Neurophysiology 126, 1071–1107 (2015).

21. Bystritsky, A. et al. A review of low-intensity focused ultrasound pulsation. Brain Stimulation 4, 125–136 (2011).

22. Deng, Z.-D., Lisanby, S. H. & Peterchev, A. V. Electric field depth–focality tradeoff in transcranial magnetic stimulation: Simulation comparison of 50 coil designs. Brain Stimulation 6, 1–13 (2013).

23. di Biase, L., Falato, E. & Di Lazzaro, V. Transcranial Focused Ultrasound (tFUS) and Transcranial Unfocused Ultrasound (tUS) Neuromodulation: From Theoretical Principles to Stimulation Practices. Front. Neurol. 10, 549 (2019).

24. Nitsche, M. A. et al. Shaping the Effects of Transcranial Direct Current Stimulation of the Human Motor Cortex. Journal of Neurophysiology 97, 3109–3117 (2007).

25. Hutcheon, B. & Yarom, Y. Resonance, oscillation and the intrinsic frequency preferences of neurons. Trends in Neurosciences 23, 216–222 (2000).

26. Gotman’, J. Relationships between Triggered Seizures, Spontaneous Seizures, and lnterictal Spiking in the Kindling Model of Epilepsy. 15.

27. Song, H. et al. Effects of Antiepileptic Drugs on Spontaneous Recurrent Seizures in a Novel Model of Extended Hippocampal Kindling in Mice. Front. Pharmacol. 9, 451 (2018).

28. Ash, C., Dubec, M., Donne, K. & Bashford, T. Effect of wavelength and beam width on penetration in light-tissue interaction using computational methods. Lasers Med Sci 32, 1909–1918 (2017).

